# Haptic assistance that restricts use of redundant solutions is detrimental to motor learning

**DOI:** 10.1101/819771

**Authors:** Rakshith Lokesh, Rajiv Ranganathan

## Abstract

Understanding the use of haptic assistance to facilitate motor learning is a critical issue, especially in the context of tasks requiring control of motor variability. However, the question of how haptic assistance should be designed in tasks with redundancy, where multiple solutions are available, is currently unknown. Here we examined the effect of haptic assistance that either allowed or restricted the use of redundant solutions on the learning of a bimanual steering task. 60 college-aged participants practiced steered a single cursor placed in between their hands along a smooth W-shaped track of a certain width as quickly as possible. Haptic assistance was either applied at the ‘task’ level using a force channel that only constrained the cursor to the track, allowing for the use of different hand trajectories, or (ii) the ‘individual effector’ level using a force channel that constrained each hand to a specific trajectory. In addition, we also examined the effect of ‘fading’ – i.e., decreasing assistance with practice to reduce dependence on haptic assistance. Results showed all groups improved with practice - however, groups with haptic assistance at the individual effector level performed worse than those at the task level. Moreover, fading of assistance did not offer learning benefits over constant assistance. Overall, the results suggest that haptic assistance is not effective for motor learning when it restricts the use of redundant solutions.

## I. INTRODUCTION

Robotic training is widely adopted to assist in the learning of novel motor tasks, especially those requiring precision. For example, a stroke survivor attempting to place a cup of coffee on a narrow ledge is faced with a task of moving the cup in a specified trajectory while controlling task variability – i.e., variability that affects the movement of the cup. Although several different algorithms have been used to explore how haptic feedback can be used to influence motor learning in such contexts [1]–[5], here we focus on ‘haptic assistance’ which is designed to minimize errors during training.

A critical issue in this regard is how to design haptic assistance to best control task variability. Prior studies have almost exclusively used non-redundant tasks where task variability can only be controlled directly by controlling the movement variability of the end-effector, i.e. enforcing the same movement from trial to trial [6]–[9]. However, when tasks have multiple degrees of freedom, the redundancy associated with this arrangement leads to a situation where task variability can be controlled without necessarily repeating the same movements at all the individual effectors. This strategy of ‘repetition without repetition’ (i.e. achieving the same task goal without repeating the same movements) has been observed extensively in human motor control [10]–[14]. However, the question of how haptic assistance has to be provided in such redundant tasks to enhance learning is not known.

Haptic assistance can be provided at two levels in redundant tasks - (i) the ‘task’ level where the assistance constrains deviations only when they interfere with the task, or (ii) the ‘individual effector’ level where the assistance constrains deviations of individual effector motions. The key distinction between these two levels is that haptic assistance at the task level allows the use of multiple redundant solutions and flexibility in movements from trial-to-trial [15]. On the other hand, haptic assistance at the individual effector level limits such flexibility from trial-to-trial, but may still be able to facilitate learning through a ‘use-dependent’ learning mechanism [16], [17].

A second issue when providing haptic assistance is that of ‘fading’ assistance. Learners with constant haptic assistance throughout practice tend to become dependent on it [18] leading to a significant deterioration in performance upon removal of assistance [19], [20]. One strategy to counter this overreliance on haptic feedback is by fading assistance– i.e. gradually decreasing assistance with practice [21]–[23], Fading can also implicitly be built into the task by implementing ‘assist-as-needed’ protocols, wherein haptic assistance is provided only outside a bandwidth of errors and the forces are increased proportionally to errors [24]. However, how the effect of fading interacts with the level of haptic assistance (i.e. task or individual effector) is not known.

Here, we examined the role of haptic assistance in learning redundant tasks. We developed a task where participants had to trace a complex trajectory using a cursor. Critically, the cursor was placed at the mean position of the two hands, which made the task kinematically redundant because the same cursor position could be achieved by different positions of the hands. We examined two specific questions in this context - (i) how does the level at which haptic assistance is provided – i.e. task or individual effector, influence motor learning, and (ii) how does the strength of haptic assistance– i.e. constant or faded, influence motor learning.

## II. METHODS

### A. Participants

60 heathy college-aged adults (age range: 18-24 years, 20 men, 40 women) participated in the study and received extra course credit for participation. All participants provided informed consent and the procedures were approved by the Institutional Review Board at Michigan State University.

### B. Apparatus

We used a bimanual manipulandum (KINARM Endpoint Lab, BKIN Technologies, ON), which consisted of two separate robotic arms that allowed motion in a 2-D horizontal plane. Each robotic arm had a handle located at the end which could be grasped by participants. Participants were seated on a height-adjustable chair and looked into a screen at around 45-degree angle below eye level as shown in Fig. 1a. The visual information was presented in such a way that the objects on the screen appear to be located in the plane of the hands. Kinematic data from both handles were sampled at 1000 Hz.

**Fig. 1.**
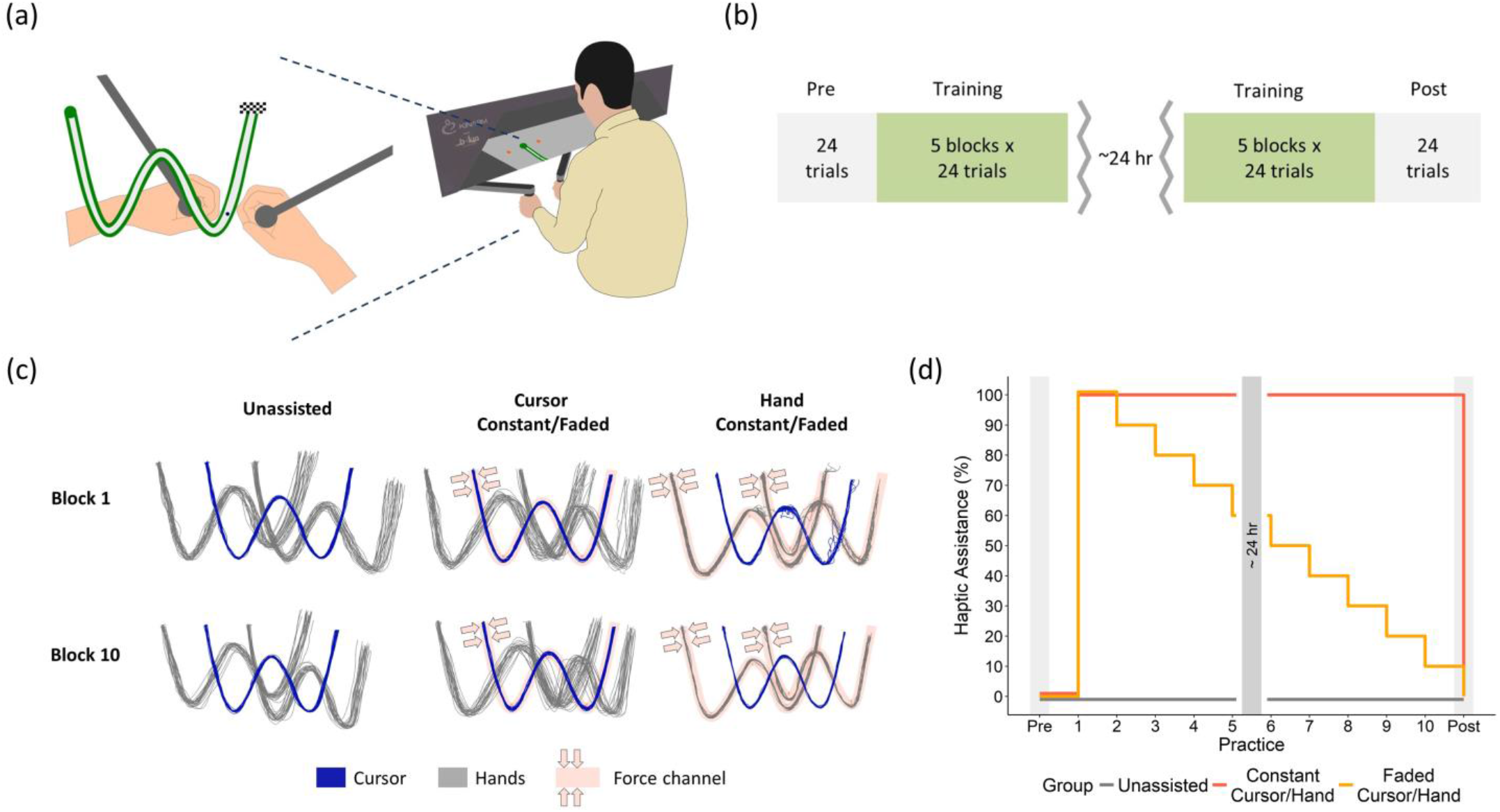
(a) Experimental setup - Participants held the handles of a bimanual manipulandum and looked at a screen that appeared to be in the plane of their hands. (left) They traced a ‘W’ shaped track using a blue cursor placed in between their hands, and the goal was to move as fast as possible while maintaining the cursor within the grey track. (b) Experimental protocol for all 5 groups (Cursor Constant, Cursor Faded, Hand Constant, Hand Faded, Unassisted). Participants did a Pre-test followed by five blocks of training on the first day, and 5 blocks of training followed by a Post-test on the second day (c) Haptic assistance using spring-like forces were applied based on cursor motion for the Cursor groups and based on individual hand motion for the Hand groups. Cursor, left hand and right hand trajectories from a representative participant from each group are shown for Block 1 and 10 in training. (d) Fading of haptic assistance. During the training blocks, Constant groups received 100% assistance, whereas the Faded groups received a linear decrease in the assistance at the start of each block. The Unassisted group did not receive any haptic assistance during training. There was no haptic assistance during the Pre-test and Post-test blocks for all groups.

### C. Task Description

The participants performed a bimanual steering task [25]. Participants controlled a cursor of diameter 4 mm and steered it from start position to end position along a smooth W-shaped track of length 738 mm (Fig. 1a). The goal of the task was to complete the movement as fast as possible, while maintaining the cursor within the grey track. The width of the track was always visible to the participant and consisted of two regions highlighted in different colors. The width of the inner grey track was 6 mm (the ‘allowed region’) and the width of the surrounding green track was 3mm. When the cursor deviated from the track, the surrounding track changed color to red serving as a visual cue to help maintain the cursor within the track.

### D. Cursor Mapping

The position of the cursor (X_C_, Y_C_) was displayed at the average position of the two hand locations, making the task redundant. This 4-to-2-mapping can be represented as shown in (1):

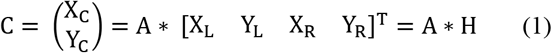

Where C is the cursor position, A is the ‘mapping matrix’ and H is the vector of the left hand and right hand coordinates.

### E. Procedures

At the start of each trial, participants saw two individual cursors (one for each hand), which allowed them to position each hand in its own start circle – this was done to ensure that the two hands always started at the same position every trial. Once each hand reached its start position, the individual cursors disappeared and were replaced by a single cursor at the average position of the two hands. Participants then moved this cursor towards the finish position as fast as possible staying within the width of the track.

To encourage participants to go faster while staying inside the track, participants were shown a score at the end of the trial. Participants started with a maximum of 100 points at the beginning of a trial and received a penalty in proportion to the time they took to complete the whole movement (t_m_) and the time that the cursor spent outside the track (t_o_) according to (2). If the cursor completely went outside the surrounding track, they were awarded zero points on that trial. In addition to the trial score, the sum of trial scores from the completed trials in the ongoing block was shown to the participants after each trial.

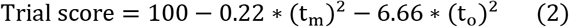

### F. Groups and Experimental Protocol

Participants were randomly assigned into 5 groups (n = 12/group) based on the mode of haptic assistance. Four groups received haptic assistance during training, and the fifth group received no haptic assistance. The four groups that received haptic assistance varied based on two factors – (i) the level at which haptic assistance was provided – at the task level (i.e. based on motion of the cursor), or at the individual effector level (i.e. based on motion of the individual hands), and (ii) the strength of the haptic assistance – constant or faded. Thus the four groups were (i) constant haptic assistance applied to the cursor (Cursor Constant – ‘CursConst’) (ii) faded haptic assistance applied to the cursor (Cursor Faded – ‘CursFade’) (iii) constant haptic assistance applied to each hand (Hands Constant – ‘HandConst’) (iv) faded haptic assistance applied to each hand (Hands Faded – ‘HandFade’). The fifth group (‘Unassisted’) did not receive any haptic assistance during training.

The experimental protocol is shown in Fig. 1b. All participants practiced initially for 10 trials without assistance, where they familiarized themselves with the task and the scoring system. After familiarization, they performed a Pre-test in which no haptic assistance was provided. This was followed by ten blocks of training spread over 2 days where each participant received haptic assistance based on their group membership. At the end of the training, participants performed a Post-test in which no haptic assistance was provided. All blocks (Pre-test, training and Post-test) consisted of 24 trials each.

### G. Haptic Assistance

Haptic assistance was provided either at the task level (i.e. based on motion of the cursor) or the individual effector level (i.e. based on the motion of the individual hands). In both cases, a compliant force field channel modelled by a spring of stiffness (K = 1 N/mm) was programmed into the task in the form of a virtual fixture. The channel applied a force (F) proportional to the deviation of the cursor/hand (Δd) from the centerline of its track in a direction perpendicular to the track according to (3). The ‘w’ here represents the width of the track, and the force was 0 as long as the cursor/hand was within the track width.

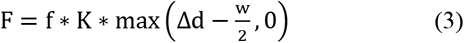

Depending on the level at which haptic assistance was introduced (task or individual effector), the channel was applied to the motion of the cursor or the two hands as shown in Fig. 1c. For the Cursor groups, the computed force was applied equally to both hands such that it appeared to have been fed through the cursor. For the Hand groups, we first obtained reference channels for each hand using the average of the Post-test hand trajectories from the participants in the Unassisted group. Each hand then felt forces independent of the other hand, based on the deviation from its own channel.

The strength of haptic assistance was either maintained constant or faded with practice in the training blocks according to Fig. 1d. We used a force factor (f) according to (3), to fade the level of haptic assistance, wherein a force factor of 2 represented the maximum haptic assistance (i.e. 100%), and a force factor of 0 represented no haptic assistance (0%).

## III. DATA ANALYSIS

### A. Block Score

The score provided to the participant on each trial was computed using (2). This score was averaged across all trials in a block for each participant.

### B. Movement Time

Movement time was defined as the time between the instant when the participant moved the cursor out of the start circle and the instant when the cursor moved into the finish box. Movement times were averaged across all trials in a block for each participant.

### C. Out of Track Time

Out-of-track time was defined as the time that the cursor was outside the track from the start to end of movement. The out of track time was then averaged across all trials in a block for each participant.

### D. Task and Null Space Variability

Since the task was kinematically redundant, the variability in hand positions was decomposed into task and null space variabilities [26]–[28]. The task space variability refers to the component of the movement variability that affects cursor motion whereas the null space variability refers to the component of the overall movement variability that has no effect on cursor motion. The path from each trial was divided into 51 spatially equidistant points from the start to the end. At each point, the corresponding hand positions from all trials in that block were extracted into a matrix H as shown in (1) and the Moore-Penrose inverse was used to decompose the hand positions into null space (H_n_) and task space (H_t_) components [25]–[27] as shown in (4) and (5) respectively, where I_4_ is an identity matrix of size 4.

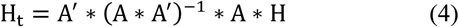

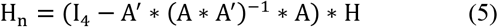

The variances of the null and task components of the hand positions were computed and summed to obtain null space and task space variability in each block.

### E. Haptic Force Reliance

Because the haptic forces that participants experienced depended both on the error as well as the time they spent outside the track, the haptic reliance on each trial was calculated by computing the net force impulse – i.e. integrating the forces experienced by the participant from start to end of movement. Note that the haptic reliance was zero for the Unassisted group during training, and in the Pre-test and Post-test block for all groups since there was no haptic assistance provided in these cases.

## IV. STATISTICAL ANALYSIS

Our primary research questions were to determine the effect of the level of haptic assistance - HapticLevel (cursor/hand) and the strength of haptic assistance - HapticStrength (constant/faded) on the task outcome variables. Because the block score, movement time and the out-of-track time are mathematically related according to (2), we show all three variables on the graphs, but exclude the out-of-track time from the statistical analysis.

### Training phase

To examine if our manipulations had the desired effect during training, we tested for three specific effects: (i) Effect of haptic assistance on performance, i.e. whether haptic assistance enhanced performance relative to no assistance, (ii) Effect of level of haptic assistance on variability, i.e., whether the Hand groups have lower null space variability relative to the Cursor groups (indicating less use of redundancy), and (iii) Effect of haptic assistance on force reliance, i.e. whether the Faded groups showed less reliance on haptic assistance relative to the Constant groups. All these tests were examined by using a 2×2 (HapticLevel x Haptic Strength) ANOVA on the last block of training. Bonferroni adjusted contrasts were used for post-hoc analysis and to make comparisons with the Unassisted group.

### Test phase

To examine the effect of learning in groups that received haptic assistance, we only used the test phases (i.e. pre- and post-test). Because our groups were based on a 2 × 2 design (HapticLevel × Haptic Strength), we used a 2 × 2 ANCOVA on the Post-test values with Pre-test values as covariate, and HapticLevel and HapticStrength as factors.

Finally, to compare the effects of haptic assistance relative to no haptic assistance, we used Bonferroni adjusted contrasts with respect to the Unassisted group on the Post-test. The significance level for all tests was set at α = 0.05.

## V. RESULTS

To examine any outliers, we compared the overall change in the Block score from the pre- to post-test for all groups. Using Tukey’s outlier criterion (i.e. above 1.5 IQR of the third quartile or below 1.5 IQR of the first quartile), we eliminated two participants from further statistical analysis (one from HandConst and one from HandFade).

### A. Training Phase

Effect of haptic assistance on performance. We found two effects of haptic assistance - the haptic groups had higher scores relative to the Unassisted group, and the constant groups had higher scores relative to the faded groups (Fig. 2a). The ANOVA on Block 10 revealed a significant effect of HapticStrength (F(1,42) = 59.74, p<0.001) and no significant effect of HapticLevel (F(1,42) = 1.96, p=0.16). Bonferroni adjusted contrasts on Block 10 showed that Constant groups had higher scores than the Faded groups (p<0.001), and both the Faded and Constant groups had higher scores than the Unassisted (p<0.001), group.

**Fig. 2.**
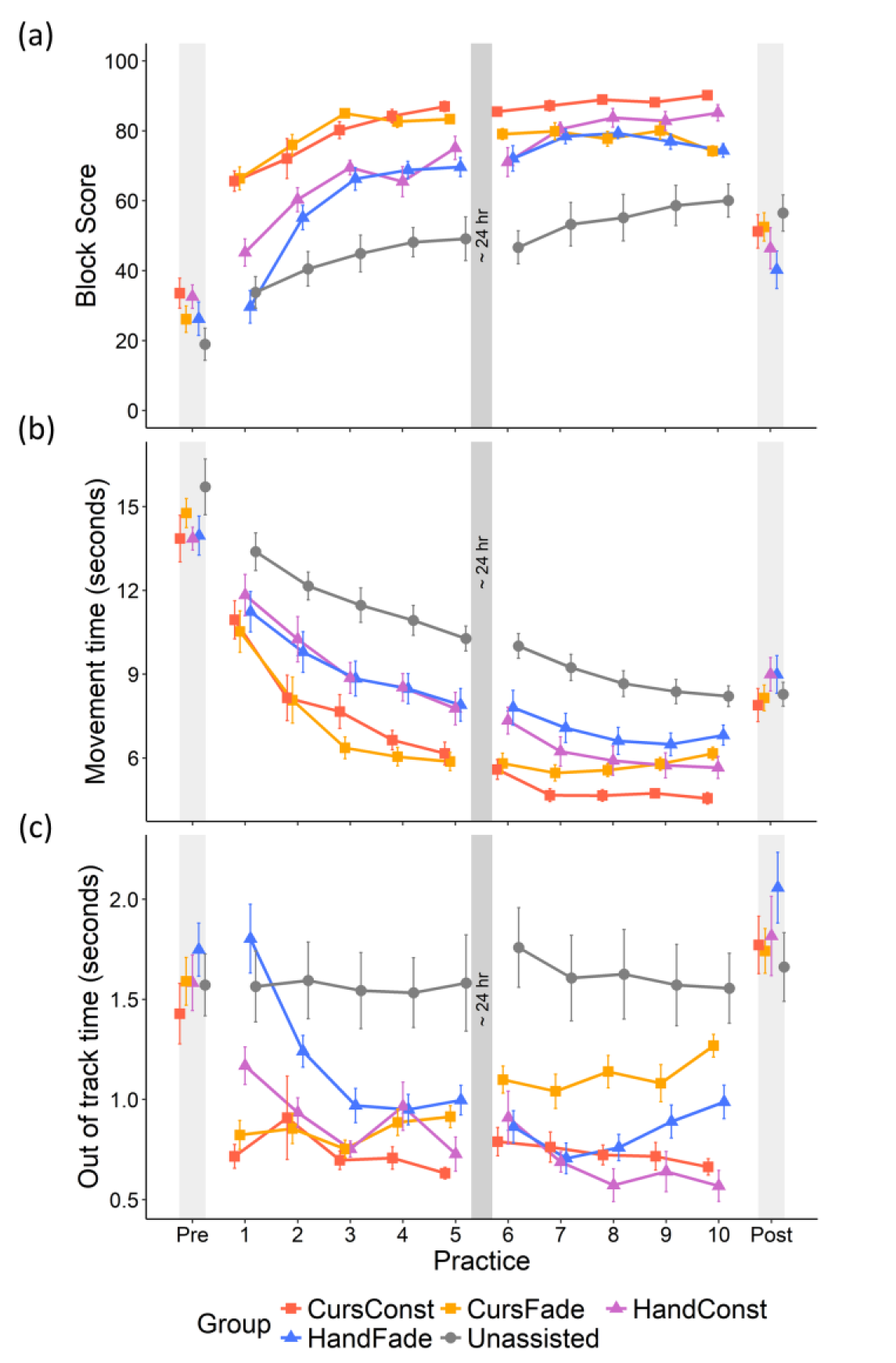
Plots of performance variables versus practice. (a) Block score- All groups improved scores with practice, but the Hand groups had relatively lower mean scores compared to the Cursor groups and the Null group (b) Movement time- All groups showed decreasing movement time with practice, and the Hand groups had relatively higher mean movement times in comparison to the Cursor and Null groups in the Post-test (c) Out of track time- Out of track times remained similar from Pre to Post, and the Hand groups showed relatively higher mean out of track times in comparison to the Cursor groups and the Null group in the Post-test.

**Fig. 4.**
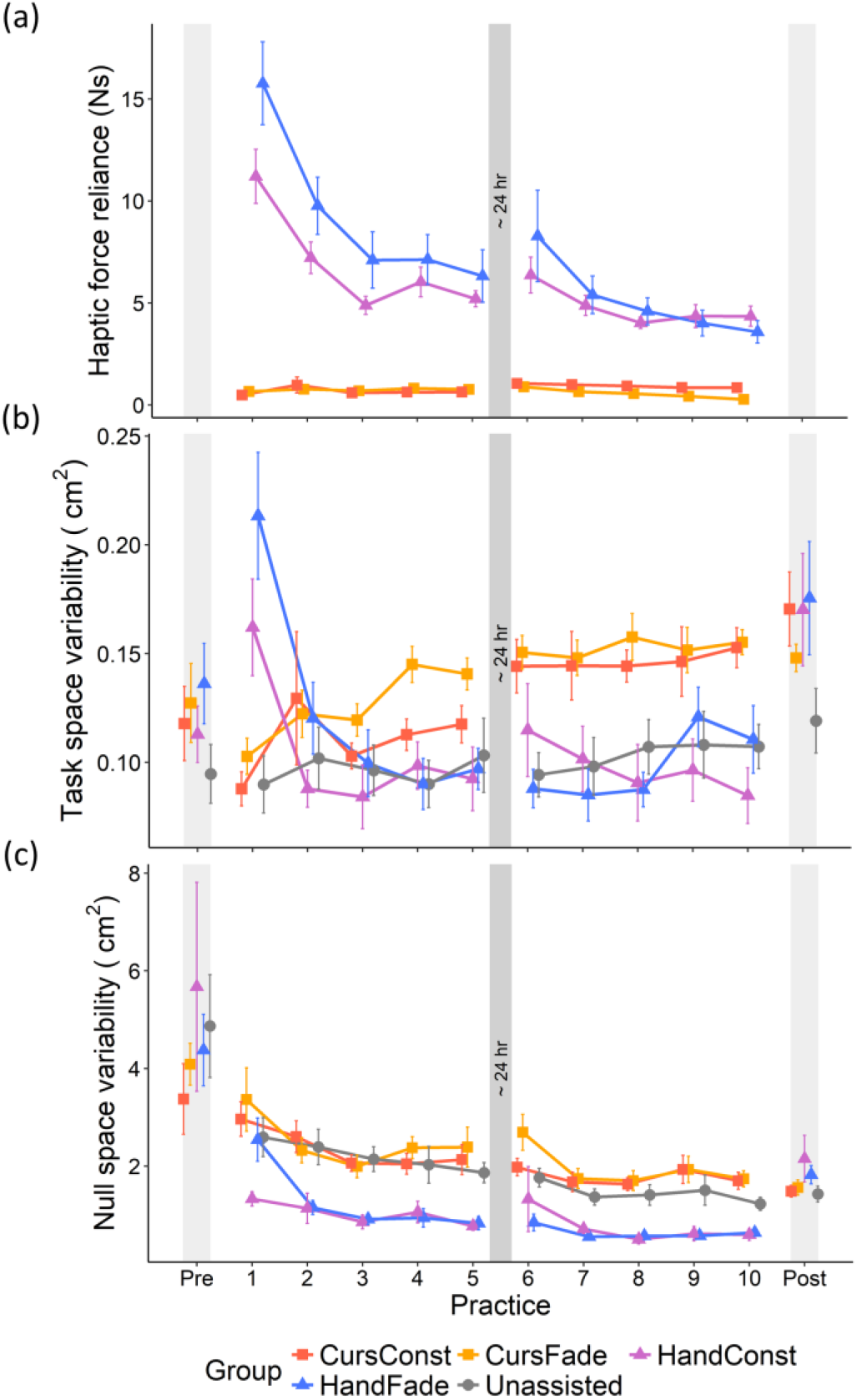
Plots of computed variables versus practice. (a) Haptic force reliance- The hand groups experienced greater but reducing amounts of haptic force in training in comparison to the Cursor groups (b) Task space variability- The Cursor groups showed increasing task space variability whereas the other groups showed reducing or unchanging task space variability with practice (c) Null space variability- Null space variability reduced with practice for all groups, but due to our haptic manipulation the Hand groups had lower null space variability in comparison to the Cursor groups in training.

Effect of haptic assistance on variability. As expected, the Hand groups had lower null space variability in comparison to Cursor and Unassisted groups, indicating reduced use of redundant solutions (Fig. 3c). The ANOVA on Block 10 revealed a significant effect of HapticLevel (F(1,42) = 60.89, p<0.001) and no significant effect of HapticStrength (F(1,42) = 0.081, p=0.77). Bonferroni adjusted contrasts showed that the Hand groups had lower null space variability relative to the Cursor groups (p<0.001). Moreover, the Hand groups had lower null space variability relative to the Unassisted group (p=0.0018), whereas the Cursor groups had higher null space variability relative to the Unassisted group (p=0.012).

Effect of haptic assistance on force reliance. The Faded groups showed similar force reliance as that of the Constant groups within each HapticLevel factor, whereas the Hand groups experienced greater force reliance than the Cursor groups (Fig. 3a). The ANOVA on Block 10 revealed a significant effect of HapticLevel (F(1,42) = 92.16, p<0.001) but no significant effect of HapticStrength (F(1,42) = 3.61, p=0.064).

### B. Test Phase

#### 1) Block Score

The Cursor groups had higher scores relative to the Hand groups in the Post-test relative to Pre-test scores (Fig. 2a). The ANCOVA indicated a significant effect of HapticLevel (F(1,41)=4.31, p=0.044), and no other significant effects.

Comparison to Unassisted group. Bonferroni adjusted contrasts in the Post-test revealed no significant differences between Hand and Unassisted (p=0.071), and Cursor and Unassisted (p=0.89).

#### 2) Movement Time

The Hand groups had higher movement times in comparison to the Cursor groups in the Post-test with respect to the Pre-test times (Fig. 2b). The ANCOVA indicated a significant effect of HapticLevel (F(1,41)=5.48, p=0.024), and no other significant effects.

Comparison to Unassisted group. Bonferroni adjusted contrasts in the Post-test revealed no significant differences between Hand and Unassisted (p=0.58), and Cursor and Unassisted groups (p=0.99).

#### 3) Task Space Variability

The Hand and Cursor groups had similar task space variabilities in the Post-test with respect to Pre-test variabilities (Fig. 3b). The ANCOVA indicated no significant effect of HapticLevel or HapticStrength or their interaction.

Comparison to Unassisted group – Bonferroni adjusted contrasts on the Post-test revealed higher task variabilities in the Hand group relative to the Unassisted group (p=0.044), but no significant difference between Cursor and Unassisted groups (p=0.16).

#### 4) Null Space Variability

The Hand and Cursor groups had similar null space variabilities in the Post-test with respect to the Pre-test variabilities (Fig. 3c). There was no significant effect of HapticLevel or HapticStrength or their interaction.

Comparison to Unassisted group. Bonferroni adjusted contrasts in the Post-test revealed no significant differences between Hand and Unassisted (p=0.133), and Cursor and Unassisted (p=0.99).

## VI. DISCUSSION

The goal of this study was to examine haptic assistance in the learning of tasks with redundancy. We specifically asked two questions - (i) how does the level at which haptic assistance is provided – i.e. task or individual effector, influence motor learning, and (ii) how does the strength of haptic assistance – i.e. constant or faded influence motor learning. We found that (i) haptic assistance at the individual effector level was detrimental to motor learning relative to the task level, and (ii) fading haptic assistance had no beneficial effect on learning relative to constant haptic assistance in our context.

When we examined the overall amount of learning based on the level of haptic assistance (task or individual effector), we found that all groups improved their performance substantially from pre- to post- test (movement times were cut by almost ~40% from pre- to post- test). However, the groups that received assistance at the level of individual effectors (i.e. the Hand groups) performed worse compared to the groups that received assistance at the task level (i.e. the Cursor groups). This was mainly driven by changes in movement time, with the Cursor groups going faster than the Hand groups. One potential reason for this effect is that the Hand groups had limited use of redundancy as evidenced by the lower null space variability during training. This meant that participants in these groups were not able to use the redundancy in the task to flexibly change their individual hand trajectories from trial to trial. Moreover, the use of redundant solutions also seemed to be a ‘natural’ tendency for the nervous system, which was impaired in the Hand groups. This was reflected by the increased reliance on haptic forces in training and the sudden increase in null space variability during the post-test when the haptic forces were removed.

These results are consistent with theoretical perspectives [13] such as the uncontrolled manifold [10], [29] and optimal feedback control [11], [30] which suggest a critical role for the ‘null space’ in these redundant tasks. One particular idea is that the null space acts as a ‘noise buffer’ allowing task variability to be small; as a result, controlling the null space variability might have had a negative effect on learning the task. Prior studies in multi-effector coordination tasks typically have shown that practicing with individual effectors sequentially is less effective than practicing simultaneously with the available redundancy [31]. Here, we further strengthen this argument by showing that even when groups perform simultaneous bimanual movements, the group that is restricted in its use of redundant solutions shows poorer learning.

When comparing the groups that received haptic assistance with the Unassisted group, we found that in general, no group outperformed the Unassisted group. Even though the haptic groups had better performance over the unassisted group in the training blocks, they could not retain same levels of performance in the post-test when the haptic assistance was removed. These results are consistent with prior work showing that haptic assistance has a stronger influence on performance but did not enhance learning [20]. While these results support the ‘specificity of practice’ principle [32], [33] (i.e. that learning is best when training conditions match testing conditions), it is important to note that the haptic assistance groups (esp. the Cursor groups) also did not perform significantly worse than the unassisted group. This indicates that haptic assistance may be especially useful in contexts where it may not be feasible to experience large errors even during training (for e.g., if there are safety issues involved with experiencing large errors) [21]

Finally, with respect to the effect of fading, surprisingly we found no significant effects of fading on learning. There are two possible reasons for this – first, because assistance was only applied when the cursor or hand exceeded the channel boundary, as participants performed better on the task, this naturally leads to a decrease in the reliance on haptic assistance, even though the strength of the haptic assistance was not changed. Second, the fading of the assistance was done in an open-loop fashion (i.e. all participants got the same strength regardless of performance) and may not have been optimal in our case because participants may not have had enough practice at a given haptic strength before moving to the next lower strength level. This is supported by the observation that even the Faded groups experienced a significant drop in performance going from the training block to the post-test. We speculate that fading could be more effective if it is made ‘closed-loop’ and tied to task performance by using performance adaptive assistance algorithms [19], [34]–[36].

The current results potentially have important implications for the design of robots for rehabilitation. With the rise in the use of exoskeletons for learning and rehabilitation, a big unanswered question is how these devices need to be used to facilitate learning. Previous results have suggested that strategies that allow some degree of variability are important for motor learning [37], [38]. Our results here further add to this evidence by showing that not only is variability important, but preserving the ability of the nervous system to use redundant solutions during learning is critical for learning. Therefore, rather than enforcing a ‘single’ movement pattern, it is likely that exoskeletons that allow for the use of these redundant solutions would be optimal for rehabilitation.

